# Typhi Mykrobe: fast and accurate lineage identification and antimicrobial resistance genotyping directly from sequence reads for the typhoid fever agent *Salmonella* Typhi

**DOI:** 10.1101/2024.09.30.613582

**Authors:** Danielle J. Ingle, Jane Hawkey, Martin Hunt, Zamin Iqbal, Jacqueline A. Keane, Ayorinde O. Afolayan, Niyaz Ahmed, Saadia Andleeb, Philip M. Ashton, Isaac I. Bogoch, Megan E. Carey, Marie Anne Chattaway, John A. Crump, Paula Diaz Guevara, Benjamin P. Howden, Hidemasa Izumiya, Jobin John Jacob, Louise M. Judd, Arti Kapil, Karen H. Keddy, Justin Y. Kim, Myron M. Levine, Masatomo Morita, Satheesh Nair, Sophie Octavia, Iruka N. Okeke, Precious E. Osadebamwen, Sadia Isfat Ara Rahman, Assaf Rokney, David A. Rasko, Varun Shamanna, Michael J. Sikorski, Anthony M. Smith, Gabriel T. Sunmonu, Kaitlin A. Tagg, Ryan R. Wick, Zoe A. Dyson, Kathryn E. Holt, Global Typhoid Genomics Consortium

## Abstract

**Background:** Typhoid fever results from systemic infection with *Salmonella enterica* serovar Typhi (Typhi) and causes 10 million illnesses annually. Disease control relies on prevention (water, sanitation, and hygiene interventions or vaccination) and effective antimicrobial treatment. Antimicrobial resistant (AMR) Typhi lineages have emerged and become established in many parts of the world. Knowledge of local pathogen populations informed by genomic surveillance, including of lineages (defined by the GenoTyphi scheme) and AMR determinants, is increasingly used to inform local treatment guidelines and to inform vaccination strategy. Current tools for genotyping Typhi require multiple read alignment or assembly steps and have not been validated for analysis of data generated with Oxford Nanopore Technologies (ONT) long-read sequencing devices. Here, we introduce Typhi Mykrobe, a command line software tool for rapid genotyping of Typhi lineages, AMR determinants, and plasmid replicons direct from sequencing reads.

**Results:** We validated Typhi Mykrobe lineage genotyping by comparison with the current standard read mapping-based approach and demonstrated 99.8% concordance across nearly 13,000 genomes sequenced with Illumina platforms. For the few isolates with discordant calls, we show that Typhi Mykrobe results are better supported by the evidence from raw sequence read data than the results generated using the mapping-based approach. We also demonstrate 99.9% concordance for detection of AMR determinants compared with the current standard assembly-based approach, with similar results for plasmid marker detection. Typhi Mykrobe predicts clinical resistance categorisation (S/I/R) for eight drug classes, and we show strong agreement with phenotypic categorisations generated from reference laboratory minimum inhibitory concentration (MIC) data for n=1,572 Illumina-sequenced isolates (>99% agreement within one doubling dilution). We show strong concordance (>96% for genotype and >98% for AMR and plasmid) between calls made from ONT reads and those made from Illumina reads for isolates sequenced on both platforms (n =93 genomes). Typhi Mykrobe takes less than a minute per sample and is available at https://github.com/typhoidgenomics/genotyphi.

**Conclusions:** Typhi Mykrobe provides rapid and sensitive genotyping of Typhi genomes direct from Illumina and ONT reads, although lower accuracy was observed for R9 ONT data. It demonstrated accurate assignment of GenoTyphi lineage, detection of AMR determinants and prediction of corresponding AMR phenotypes, and identification of plasmid replicons.

## Background

Typhoid fever is caused by systemic infection with the human-restricted bacterium *Salmonella enterica* subspecies *enterica* serovar Typhi (Typhi) [1]. More than 10 million typhoid fever illnesses occur annually (mainly in South Asia), associated with 100,000 deaths, and the risk of complications and death increases without effective antimicrobial therapy [2]. The World Health Organization AWaRe (Access, Watch, Reserve) Antibiotic guidance recommends typhoid fever is treated with ciprofloxacin, unless resistance is prevalent locally, in which case azithromycin is recommended to treat uncomplicated disease and intravenous ceftriaxone is recommended for treatment of severe disease [3,4]. A high proportion of typhoid fever is caused by antimicrobial resistant (AMR) Typhi, and nearly all Typhi isolated in South Asia have been ciprofloxacin non-susceptible for over two decades [5,6]. Extensively drug resistant (XDR, resistant to amoxicillin/ampicillin, chloramphenicol, trimethoprim-sulfamethoxazole, ciprofloxacin, and ceftriaxone) Typhi, for which the only treatment options are oral azithromycin or intravenous carbapenems, have been circulating in Pakistan for nearly a decade with travel-associated infections reported globally [7–9]. Concerningly, XDR Typhi with no travel links has been reported in USA and China [7–10], and XDR isolates with additional resistance to azithromycin and carbapenems were recently reported in Pakistan [11].

Tracking the emergence and spread of AMR Typhi through whole genome sequencing (WGS) is important to inform public health control measures such as deployment of typhoid conjugate vaccines (TCVs) [12], investigating and controlling outbreaks [10], and informing the development of antimicrobial treatment guidance [3], including management of travel-associated infections in non-endemic countries [4]. Genetic determinants of AMR are well-understood in Typhi [13,14], and include both acquired genes that are plasmid-borne or chromosomally integrated [15], and point mutations in core genes. These include *acrB* for azithromycin resistance [16], and quinolone resistance-determining regions (QRDR) of *gyrA*, *gyrB,* and *parC* for ciprofloxacin non-susceptibility and resistance [13,17]. Recently, carbapenem-resistant Typhi was described, due to acquisition of a plasmid-borne carbapenemase gene [11]. Typhi is highly clonal, which has hampered subtyping in the past; however, its population structure has been well defined using WGS [15], and the GenoTyphi framework for genotyping and variant nomenclature first proposed in 2016 [18] has been widely adopted by the global community to facilitate identification of emerging variants and communication across laboratories and settings [13,14,19–21].

The GenoTyphi scheme was originally defined based on nearly 2,000 WGS isolates from over 60 countries [18]. It uses a hierarchical framework based on the global phylogeny, stratifying the Typhi population structure into four primary lineages, 16 clades, and 49 subclades (henceforth referred to as genotypes), based on lineage-specific marker single nucleotide variants (SNVs). A schematic mapping the hierarchical genotype labels to the Typhi phylogeny is shown in **Supplementary Figure 1**. Under the GenoTyphi scheme, the lineage formerly known as ‘H58’ was designated genotype 4.3.1, and is further divided into multiple higher-resolution genotypes with the prefix 4.3.1 including 4.3.1.1, 4.3.1.2 (see **Supplementary Figure 1**). The H58 lineage, or genotype 4.3.1 and associated sub-genotypes, has been associated with the spread of multidrug resistance (MDR, resistant to amoxicillin/ampicillin, chloramphenicol, trimethoprim-sulfamethoxazole) Typhi throughout Asia and into Eastern Africa [15]. The scheme was later extended to provide further discriminatory power for identifying additional subpopulations of epidemiological importance [21–27], such as the XDR Typhi lineage associated with outbreaks in Pakistan (genotype 4.3.1.1.P1) [26,27]. The scheme currently includes 87 genotypes [28], and its future maintenance and development will be managed by a working group within the Global Typhoid Genomics Consortium (https://www.typhoidgenomics.org/).

The GenoTyphi scheme specification consists of a set of marker SNVs defining the hierarchical genotypes, which in principle can be used by any bioinformatics tool to assign genotypes to new genomes. The original GenoTyphi paper [18] was accompanied by a Python-based pipeline (available at: https://github.com/typhoidgenomics/genotyphi) designed for typing Illumina WGS data against the scheme. This implementation requires users to first map short reads or assemblies to the reference sequence of Typhi strain CT18 (accession AL513382.1) [29], then provide as input the resulting binary alignment map (BAM) or variant call file (VCF). Marker SNVs were identified from these input files and used to assign genomes to genotypes.

Functionality was extended to include detection of mutations associated with reduced susceptibility to ciprofloxacin or azithromycin [22]. This mapping-based approach had various limitations, largely due to the need for users to pre-map their data to a specific reference, which was cumbersome (e.g. there is wide variation in the upstream mapping tools used and some users had issues with mapping tools creating incompatible output files, the BAM files are large and inefficient to process), and AMR analysis was restricted to mutation detection. The GenoTyphi scheme has also been implemented in other genotyping tools, including Typhi PathogenWatch [13], which works by analysing assembled genomes, and the split-*k*-mer-based BioHansel tool, which does not include AMR detection [30].

Here, we present a novel approach to rapid genotyping of both AMR and GenoTyphi lineages from reads, implemented in the Mykrobe open-source software framework [31], dubbed Typhi Mykrobe. Input into this new tool accepts raw WGS reads (FASTQ files), and has been tested with reads generated by ONT and Illumina platforms. The tool performs rapid (i) lineage genotyping using the GenoTyphi framework; (ii) detection of a broad range of clinically relevant molecular determinants of AMR, including both acquired genes and mutations; and (iii) detection of plasmid replicon markers common to Typhi. We validated GenoTyphi lineage assignment from reads by comparison with the original mapping-based implementation, using 12,839 Typhi genomes [14]. We assessed AMR genotype calls by comparison with assembly-based genotype calls from PathogenWatch, and with antimicrobial susceptibility phenotypes for 4,018 isolates. Finally, we assessed performance on long read data generated using Oxford Nanopore Technologies (ONT) MinION devices, using isolates with matched ONT and Illumina data.

### Implementation

As outlined in the Introduction, GenoTyphi is a typing scheme specific to Typhi, which specifies marker SNVs suitable to identify specific lineages. In principle GenoTyphi can be used with any genotyping tool that is capable of searching for the marker SNVs in sequence data. GenoTyphi provides the framework directing genotyping tools what SNVs to look for and how to interpret them in terms of lineages to report. Mykrobe is a genotyping tool, which uses a *k*-mer-based approach to identify marker SNVs in sequence reads. It was developed for typing *Mycobacterium tuberculosis* and *Staphylococcus aureus* [32–34], but can in principle be used with any genotyping scheme. Here we present a ready-to-run code base that allows users to type genomes from raw reads, using Mykrobe, against the GenoTyphi framework to identify and report known lineages – which we label “Typhi Mykrobe”. This tool additionally includes AMR and plasmid typing frameworks, and generates predictions of resistance to the drugs recommended for clinical treatment of typhoid. Details of this Implementation are described below.

### Genotyping targets

The current scope of the GenoTyphi scheme as discussed here (n=87 genotypes) is illustrated in **Supplementary Figure 1**. This includes five new genotypes (3.5.4.1, 3.5.4.2, 3.5.4.3, 4.3.1.2.1, 4.3.1.2.1.1) recently described in a technical report [28]. The full scheme specification is available in the GenoTyphi repository (https://github.com/typhoidgenomics/genotyphi) in the file Genotype_specification.csv.

The AMR and plasmid targets included in Typhi Mykrobe are summarised in **Figure 1**. These include sequence probes targeted to detect presence/absence of epidemiologically relevant mobile AMR determinants [13,14], conferring resistance to the currently recommended drugs ciprofloxacin, ceftriaxone, and azithromycin, as well as older drugs ampicillin, chloramphenicol, trimethoprim-sulfamethoxazole, and tetracycline, as well as carbapenems. Fourteen SNVs associated with resistance to ciprofloxacin and azithromycin are also included. In addition, Typhi Mykrobe includes probes to detect the presence or absence of 13 plasmid replicons previously reported in Typhi [14], including 11 associated with AMR plasmids, the cryptic plasmid pHCM2, and linear plasmid pBSSB1, which carries the alternative flagellin z66 [35]. A marker SNV to identify the plasmid sequence type 6 (PST6) lineage of the dominant IncHI1 plasmid is also included [36]. The full list of AMR targets is available in the GenoTyphi repository (https://github.com/typhoidgenomics/genotyphi) in the file AMR_genes_mutations_plasmids.csv.

**Figure 1:**
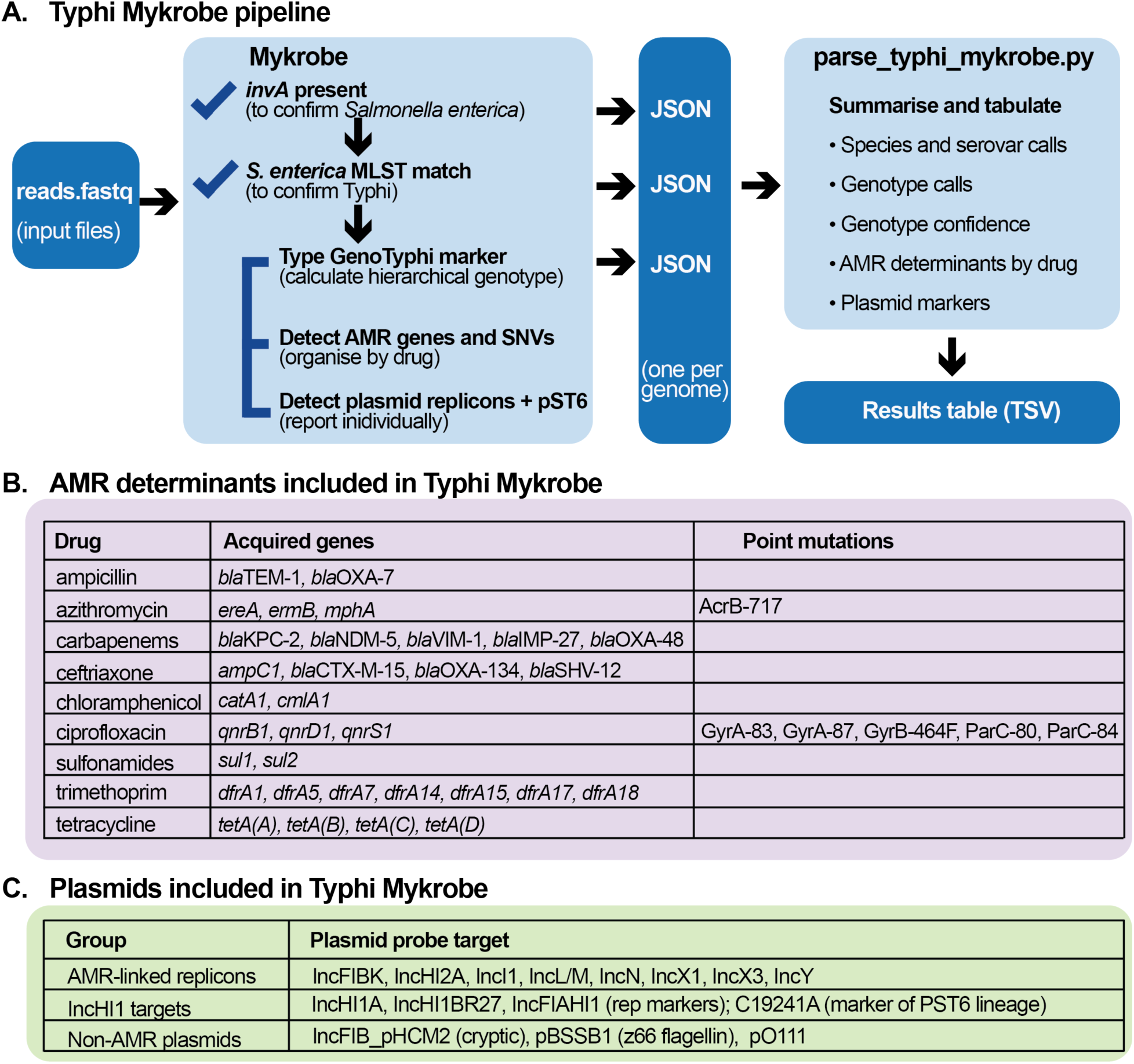
Typhi Mykrobe functionality A) A schematic overview of the Typhi Mykrobe pipeline. B) Table of epidemiologically important AMR determinants targeted for detection/typing by Typhi Mykrobe, grouped by drug. C) Table of epidemiologically important plasmid replicons targeted for detection by Typhi Mykrobe.

### Implementation in Mykrobe

The Mykrobe software, first developed in 2015 for *Mycobacterium tuberculosis* and *Staphylococcus aureus* [32], uses a *k*-mer-based approach to identify marker SNVs and the presence or absence of target genes using ‘probe’ sequences to screen against [31]. These data are then used to assign hierarchical genotypes and report AMR determinants. Probe sets have been previously developed and implemented for other bacterial pathogens including *Mycobacterium tuberculosis* [32–34], *Staphylococcus aureus* [31], *Shigella sonnei* [37], and *Salmonella* Paratyphi B [38]. Mykrobe is available at https://github.com/Mykrobe-tools/mykrobe.

Mykrobe probe sets were created for genotype and AMR marker SNVs using the *mykrobe variants make-probes* command using a *k*-mer size of 21 and the Typhi CT18 reference genome (accession: AL513382.1). The nested hierarchical relationships of the marker SNVs were specified and Mykrobe used this information to identify the best supported genotype call (e.g., a call of ‘4.3.1.1’ is typically supported by detection of nested markers that demarcate different levels of the hierarchy: 4, 4.3.1, 4.3.1.1). Briefly, probe sets for gene presence or absence were created using a Python script that extracted the relevant AMR sequences. Details of the specific commands and Python scripts are provided in https://github.com/typhoidgenomics/genotyphi. Probe sets were also created to confirm input read sets as *S. enterica* using the presence of marker gene *invA*, involved in invasion of intestinal epithelium cells by *S. enterica*, and serovar Typhi based on the 26 known multi-locus sequence types (STs) (see **Figure 1**). Further details of the preliminary testing and implementation of the Typhi probe sets are available in the technical report https://zenodo.org/doi/10.5281/zenodo.7407984. The Typhi Mykrobe probe sets and typing panel are available at https://doi.org/10.6084/m9.figshare.27102334.

### Summarising Mykrobe outputs

To support interpretation of the Mykrobe output, we developed a Python-based parser script *parse_typhi_mykrobe.py,* which takes as input a set of JavaScript Object Notation (JSON) files output by Mykrobe, and summarises them in a tab-delimited table with one row per genome, and columns to indicate the genotype calls (see **Figure 1**). Also included are several quality-control fields reported by Mykrobe, including read-level support for each marker SNV contributing to the final genotype call (‘node support’), and for any additional markers detected. This information is helpful for identifying contaminated samples (see examples in Results). The parser script also assigns a summary ‘confidence’ level in the final GenoTyphi lineage call, based on the Mykrobe quality scores for each nested component marker that makes up the final genotype call. In brief, a weak’ confidence call is assigned when one or more marker SNVs are of low quality, defined as Mykrobe quality score of 0 or Mykrobe quality score of 0.5 with minority support (i.e., <50% of reads) for the derived allele. A ‘moderate’ confidence call is assigned if one (and only one) marker SNV has a Mykrobe quality score of 0.5 but with majority support (i.e. ≥50% of reads) for the derived allele. Otherwise, the genotype call is assigned ‘strong’ confidence.

The parser script summarises the AMR markers detected by Mykrobe in the form of a pseudo-antibiogram with one column per drug. Each drug column contains either ‘S’ (indicating no markers detected) or ‘R’ (detected markers). For ciprofloxacin, it is useful to distinguish between low-level resistance (MIC 0.06-0.5 mg/L), which is associated with a single mutation or acquired gene and high-level resistance (MIC >0.5 mg/L), which is associated with combinations of two or more determinants [14,17], because while both groups are associated with increased fever clearance times and probability of treatment failure compared with susceptible strains, high-level resistance is associated with worse outcomes than low-level resistance [39]. Current guidelines and terminology for these groups vary, but for simplicity, the Mykrobe parser reports these groups as ‘I’ (for intermediate) and ‘R’, respectively. This matches the current CLSI guidelines (https://clsi.org/standards/products/microbiology/documents/m100/). However, it should be noted that (i) EUCAST defines the clinical breakpoint for ‘R’ in *Salmonella* spp. as MIC >0.06 mg/L (https://www.eucast.org/clinical_breakpoints), which includes both low-level and high-level resistance; and (ii) previous literature has referred to the low-level range as ‘decreased ciprofloxacin susceptibility’ rather than ‘I’ [40]. In addition to this antibiogram-style summary, individual AMR and plasmid markers are reported in their own columns, coded as 1=present and 0=absent (as seen in **Supplementary Table 1,** available at https://github.com/typhoidgenomics/TyphoidGenomicsConsortiumMykrobe).

All code and instructions for genotyping isolates with GenoTyphi using Typhi Mykrobe are freely available in the GitHub repository at https://github.com/typhoidgenomics/genotyphi (DOI: 10.5281/zenodo.13859721).

### Validation data

Validation on Illumina short-read data was conducted using the Global Typhoid Genomics Consortium dataset described in Carey et al [14]. Reads were mapped to the CT18 reference genome (accession: AL513382) [29] using bwa-mem v0.7.17 via the Centre for Genomic Pathogen Surveillance (CGPS) mapping pipeline v1.2.2 (https://gitlab.com/cgps/ghru/pipelines/snp_phylogeny), to generate alignments in BAM format. Assemblies were generated using the CGPS assembly pipeline v2.1.0 (https://gitlab.com/cgps/ghru/pipelines/dsl2/pipelines/assembly/) which utilises the SPAdes assembler (v3.12.0) [41,42]. A total of 141 isolates were excluded due to excessive unclustered heterozygous SNVs indicative of mixed cultures (n=92), assembly issues (n=46) or both (n=3), resulting in 12,839 isolates for Mykrobe validation (listed in **Supplementary Table 2,** available at https://github.com/typhoidgenomics/TyphoidGenomicsConsortiumMykrobe). The genomes of isolates called as ‘not Typhi’ were characterised for presence of the marker genes of *invA* and the seven MLST genes using Blast in Bandage for *invA* [43] and MLST with the ‘senterica’ database (https://github.com/tseemann/mlst). The dataset covered 85 of the 87 defined genotypes (exceptions being genotypes 1 and 2.3, which represent internal nodes of the hierarchy for which no genomes have been observed). Individual calls are given in **Supplementary Table 1** and **Supplementary Table 2.**

### Validation of genotype calls

Typhi Mykrobe v0.12.1 was run on all (n=12,839) Illumina FASTQ files using the new Typhi panel (v20221208) and the results were summarised using *parse_typhi_mykrobe.py*. To validate Typhi Mykrobe’s GenoTyphi genotype calls, we compared them with those called by the original mapping-based approach. To do this, we used BAM file read alignments generated from the same n=12,839 FASTQs as described above as input to the Python script genotyphi.py (v2.0; available at https://github.com/typhoidgenomics/genotyphi) [18,22,28]. To validate the AMR and plasmid calls, we compared them to those called in corresponding genome assemblies by Pathogenwatch (available for n=11,992 genomes, assemblies for the other genomes were excluded as they did not meet Pathogenwatch quality-control criteria [13]). We downloaded the GenoTyphi, AMR, and plasmid replicon typing reports generated by Basic Local Alignment Search Tool (BLAST) searching assemblies for these genomes from Pathogenwatch on 7 March 2023 (GenoTyphi v20221208 specification for lineage genotypes). Categorical agreement (percentage of genomes where both tools agreed on the same call of presence or absence) was calculated separately for each marker, and collectively by drug (percentage of genomes where both tools agreed that at least one marker was present or that no markers were present) (**Supplementary Table 3,** available at https://github.com/typhoidgenomics/TyphoidGenomicsConsortiumMykrobe). Probes for plasmid-borne carbapenemase genes circulating in Enterobacterales (*bla*NDM-5, *bla*KPC-2, *bla*VIM-1, *bla*IMP-27, *bla*OXA-48) were added to GenoTyphi v20240407 (DOI: <u>10.5281/zenodo.13859721</u>) following the report of *bla*NDM-5 in Typhi from Pakistan [11], and validated using the genomes of the two distinct morphologic variants from that study (accessions: SRR22801766 and SRR22801806). Comparisons and statistical analyses were conducted in R v4.1.0 using tidyverse v1.3.1 [44,45].

### Benchmarking of run time for Mykrobe

We further investigated the average run time for Typhi Mykrobe to demonstrate the rapid time to a result. Here we used the short-read data from 100 Typhi genomes. These were randomly selected but included at least one genome from each genotype and a diversity of AMR and plasmid profiles. The ‘mykrobe predict’ command was run for the 100 Typhi genomes on three different computers where both the run time and RAM were recorded. The number of threads was also varied from 1 to 64 threads for the two HPC and 1 to 16 for the Mac laptop.

### Accuracy of AMR phenotype prediction

Antimicrobial susceptibility testing (AST) data were available for n=3,970 of the Illumina-sequenced isolates, sourced from three separate datasets (**Supplementary Table 4,** available at https://github.com/typhoidgenomics/TyphoidGenomicsConsortiumMykrobe). Minimum inhibitory concentration (MIC) data were available from the UK Health Security Agency (UKHSA) for n=852 isolates following EUCAST standards and interpretive criteria (v10, 2020) [46] and the US Centers for Disease Control and Prevention (US CDC) for n=720 isolates following CLSI standards and interpretive criteria (M100, 2023) [47]. Disk diffusion data following EUCAST standards and interpretive criteria (v8.0, 2018) were available for n=2,446 isolates collected and analysed from three countries as part of the Surveillance for Enteric Fever in Asia Project (SEAP), with multiple source labs contributing AST data and isolates for sequencing [6]. Typhi Mykrobe AMR calls following the S/I/R categorisation described above were compared with the phenotypic antimicrobial susceptibility S/I/R categorisation using the relevant CLSI or EUCAST standards for each data set, for the currently recommended drugs azithromycin and ciprofloxacin and ceftriaxone, and the older drugs ampicillin, chloramphenicol, trimethoprim-sulfamethoxazole, and tetracycline (note UKHSA tested amoxicillin rather than ampicillin; and SEAP did not test tetracycline). To assess concordance between genotype and phenotype, we followed the principles used for assessment of new AST devices by the US Food and Drug Administration (FDA), which considers (i) categorical agreement, (ii) major error rate (defined as the proportion of S isolates that test R with the new method [48,49]), and (iii) very major error rate (defined as the proportion of R isolates that test S with the new method) (**Supplementary Table 5,** available at https://github.com/typhoidgenomics/TyphoidGenomicsConsortiumMykrobe). The FDA standard requires >90% overall categorical agreement (within one doubling dilution), <3% major errors for each drug, and <1.5% very major errors for each drug; hence, we adopted these thresholds to assess the accuracy of the genotype-based prediction of phenotype. For drugs with high error rates, we investigated the relationship between individual resistance markers and phenotype, which we visualised using upset plots generated in R v4.1.0 using the ComplexUpset package v1.3.3 (10.5281/zenodo.3700590). For five isolates with unexplained trimethoprim-sulfamethoxazole resistance, we used nucleotide blast search of the genome assemblies to confirm the presence of wild-type chromosomal *folA* and *folP* (100% nucleotide match to reference sequences from Typhi strain Ty2) and screened the assemblies for other AMR determinants using CARD RGI v6.0.3 [50].

### Matched ONT and Illumina validation data

For 98 isolates, long-read ONT data were available in addition to the Illumina data; these included 68 isolates from Carey et al 2023 [14] and 30 novel isolates from this study. Sample-level details including accessions and technical specifications such as library, flow cell, device, basecaller, and ONT processing (including demultiplexing, trimming and filtering), are available in **Supplementary Table 6,** available at https://github.com/typhoidgenomics/TyphoidGenomicsConsortiumMykrobe. These varied between isolates including different flow cells (e.g., R9.4.1 and R10.4.1) and base calling methods. Read depth for each ONT dataset was assessed by using seqtk v1.2-r94 (https://github.com/lh3/seqtk) to determine total number of bases per read set, which was divided by the length of the CT18 reference genome (4,809,037 bases). For five genomes with read depth <30×, Kraken2 was used for the initial read-level classification of ONT data [51].

These five genomes were also aligned to the CT18 reference genome using minimap2 [52,53] to determine the read coverage of the *invA* and the seven core genes in the *Salmonella enterica* MLST scheme [54]. ONT reads were assembled for these five genomes and in addition to four genomes to resolve discrepant plasmid calls. Assembly was done using Flye v2.9.1-b1780 [55] with default parameters and yielding genome sizes of the expected 4.8 Mbp. Due to the high read depth (784×), the ONT data for SRR17299234 was subsampled prior to assembly using Filtlong v0.2.1 (https://github.com/rrwick/Filtlong) to --target_bases 480000000 (resulting in 99.81× depth). The genomes of isolates called as ‘not Typhi’ were characterised for presence of the marker genes of *invA* and the seven MLST genes using Blast in Bandage for *invA* [43] and mlst with the ‘senterica’ database (https://github.com/tseemann/mlst).

## Results

### Validation of GenoTyphi genotype calling

To validate Typhi Mykrobe for assigning hierarchical GenoTyphi genotypes, we ran it on n=12,839 Illumina readsets (covering all existing types) and compared the resulting genotypes to those called using the original mapping-based implementation. Five genomes were typed as ‘not Typhi’ by Mykrobe **(Supplementary Table 1**). Four of these genomes lacked the *invA* gene but MLST profiles were detected that were consistent with those on Pathogenwatch. In the remaining isolate, DRR071001, the *invA* gene was detected but no allele was reported for *hisD* in the MLST scheme, consistent with the MLST profiles from Pathogenwatch. Hence, these five isolates were reported as ‘not Typhi’ and not analysed further. Of the remaining n=12,834 genomes, Mykrobe called the same single genotype as the mapping-based approach for n=12,807 (99.79%). For a further n=17 genomes, Mykrobe and the mapping-based pipeline gave concordant reports of two genotypes being present, consistent with mixed samples (in the Typhi Mykrobe output this is reported as one primary genotype plus additional markers, while in the mapping-based pipeline the result is reported as a list of supported genotypes). Eight genomes belonged to genotype 3.5.3, which was nested in 3.5.4, breaking the usual hierarchical structure of the scheme. Typhi Mykrobe correctly reported these genomes as genotype 3.5.3 with support for both markers 3.5.4 and 3.5.3, as it has this nested relationship explicitly encoded in its hierarchy; whereas the mapping-based pipeline does not have this logic and incorrectly reported these genomes as genotype 3.5.4.

There were two instances of genuine disagreement in the genotype calls generated by Typhi Mykrobe as compared to the mapping-based approach. One isolate (ERR5243665) was called as genotype 2 by Mykrobe with strong confidence (no reads support for any alternative alleles or markers). It was genotyped as 0.1.3, albeit with low support, by the mapping-based approach, but the genotype 2 marker was also detected. This isolate clustered with other genotype 2 isolates in a distance-based phylogeny of genomes from the same study [6] (see tree at https://pathogen.watch/collection/0z5knw9jic9b-guevara-et-al-2021), suggesting that the Mykrobe call is correct. The other isolate (ERR2663783) was called as genotype 4.3.1.1 by Typhi Mykrobe with strong confidence. It was reported as “4.3.1.1,4.3.1.3.Bdq” (with low support) by the mapping approach; however, the support for the 4.3.1.3.Bdq marker was low. This isolate clustered closely with other 4.3.1.1 isolates from the same study [25](see tree at https://pathogen.watch/collection/b2k6dym3bdyq-tanmoy2018), which is consistent with the Mykrobe call.

We demonstrate that the run time for Typhi Mykrobe’s ‘mykrobe predict’ command on a modern computer was <1minute to complete (**Supplementary Figure 2**). The ‘mykrobe predict’ command can be run with multiple threads using the ‘--threads’/‘-t’ option. Up to ∼4 threads will increase performance. Using multiple threads on a very fast CPU, the run time for each genome was reduced to seconds (**Supplementary Figure 2**). Further, ‘mykrobe predict’ is very memory-efficient and will typically use less than 100 MB of RAM per genome.

### Validation of AMR genotyping

To assess the accuracy of Mykrobe for detecting the presence of AMR genes and mutations, we compared genotype calls from Mykrobe (called from *k-*mer analysis of reads) with those from Pathogenwatch (called from BLAST analysis of genome assemblies of the same read sets). We found very strong categorical agreement between the two methods at the individual marker level, with 99.88% of the 299,004 marker-genome combinations yielding the same call (summarised in **Table 2**, full details in **Supplementary Table 2**). High concordance was evident for all genetic markers of AMR (98.56–100%, **Table 2**), as well as plasmid replicon markers (99.74–100%, **Table 3**). The test dataset included all AMR and plasmid markers in GenoTyphi v2.0 implemented Typhi Mykrobe v0.12.1. The *ereA*, *ermB* and carbapenemases are very rarely reported in Typhi but were included in the Mykrobe panel in case they emerge.

**Table 1.** Validation of genotype calls. Summary of Typhi Mykrobe genotype calls (GenoTyphi scheme) for 12,839 Illumina readsets, compared with mapping-based genotype calls.

| **Typhi Mykrobe vs mapping-based genotype calls** | **Genomes (N)** |
| --- | --- |
| *invA*-negative, thus designated non-Typhi and not genotyped by Mykrobe | 5 |
| Single genotype, exact match | 12,807 |
| Two genotypes, exact match (reported differently) | 17 |
| Genotype 3.5.3, mismatch (reported incorrectly by mapping pipeline as 3.5.4 due to breaking hierarchy assumption) | 8 |
| Discordant result (low support reported by mapping-based pipeline, phylogenetic tree supports Mykrobe call) | 2 |

**Table 2.**
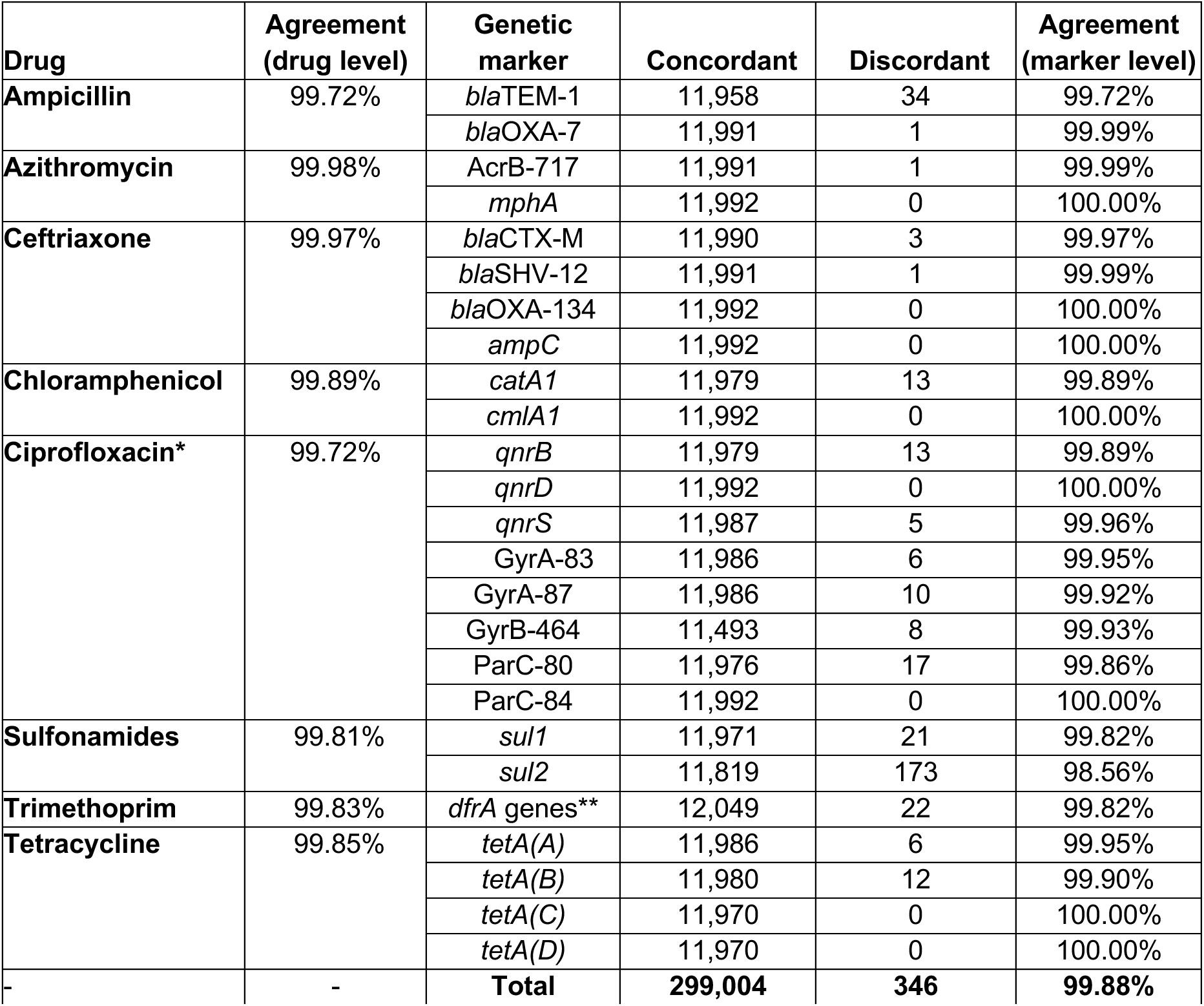
Validation of AMR genotyping (vs assembly based) Comparison of AMR genotyping by Typhi Mykrobe (*k*-mer-based analysis of reads) vs Pathogenwatch (based on nucleotide blast search of assemblies), for n=11,992 isolates. Complete details of the call comparisons are given in **Supplementary Table 3**. Categorical agreement at drug level is calculated based on agreement of the binary S/R categorisation of the 11,992 isolates by each method (i.e. ‘Concordant’ means Typhi Mykrobe and Typhi Pathogenwatch made the same call of S or R based on absence or presence, respectively, of known genetic determinants; ‘Discordant’ means one method identified determinants and called R, while the other did not and called S). *For ciprofloxacin, both tools categorise as S/I/R as defined in Methods, and categorical agreement was calculated across these three categories. **Results for *dfrA* genes are summarised together, and as some genomes have multiple *dfrA* alleles the total observations is greater than the number of isolates (see details in **Supplementary Table 3**).

**Table 3.** Validation of plasmid replicon detection (vs assembly based) . Comparison of plasmid replicon markers identified in the genomes by Typhi Mykrobe (*k*-mer-based analysis of reads) vs Pathogenwatch (PW, based on nucleotide blast search of assemblies), for n=11,992 isolates.

| **Plasmid replicon marker** | **Both** | **Mykrobe only** | **PW only** | **Agreement** |
| --- | --- | --- | --- | --- |
| IncFIAHI1 | 251 | 0 | 0 | 100% |
| IncFIB(K) | 175 | 0 | 1 | 99.99% |
| IncFIB(pHCM2) | 1406 | 26 | 5 | 99.74% |
| IncHI1A | 872 | 9 | 3 | 99.90% |
| IncHI1B | 874 | 7 | 1 | 99.93% |
| IncHI2A | 3 | 0 | 0 | 100.00% |
| IncI1 | 5 | 0 | 0 | 100.00% |
| IncL/M | 1 | 0 | 0 | 100.00% |
| IncN | 100 | 2 | 0 | 99.98% |
| IncX3 | 21 | 0 | 0 | 100.00% |
| IncY | 666 | 0 | 0 | 100.00% |

Both Mykrobe and Pathogenwatch summarise AMR data in terms of a binary phenotype categorization (S/R), assuming the presence of a marker associated with resistance to a drug implies resistance ‘R’, with the exception of ciprofloxacin, where presence of any marker is categorised as ‘I’ and specific combinations of markers are categorised as ‘R’, resulting in three categories S/I/R (see Methods). Categorical agreement was very high between the drug-level categorisations reported by both genotyping tools, with >99.7% agreement (n=2 to 33 discordant genomes) for each drug (**Table 2, Supplementary Table 3**), and for subsequent categorisation as MDR (99.86%, n=17 discordant) or XDR (99.97%, n=4 discordant).

### Validation of AMR phenotype prediction

To assess the accuracy of Mykrobe’s genotype-based clinical resistance categorisations, we compared them with AST data, which was available for a subset of n=4,018 isolates (see Methods, **Supplementary Table 4**). These isolates were from three data sets, using different methods (MIC with EUCAST standards, MIC with CLSI standards, disk diffusion with EUCAST standards), hence we analyzed the results from each dataset separately. Categorical agreement was high (>95%) for all drugs across all three datasets, and very high (>99%) for the two sets of reference-laboratory MIC measures (**Figure 2, Supplementary Figure 5**).The only exception was US CDC categorisation of low-level ciprofloxacin resistance (using the CLSI ‘I’ threshold, ≥0.125 to 0.5 mg/L), which showed 97.36% agreement with Mykrobe calls of ‘I’ or ‘R’. This was mainly due to the presence of major errors (susceptible isolates with AMR determinants detected, thus reported as R by Mykrobe, see Methods), whereby n=17 isolates tested susceptible to ciprofloxacin (n=14 with MIC=0.06 mg/L, n=3 with MIC=0.015 mg/L) but Mykrobe detected a known QRDR mutation (**Figure 3b**); categorical agreement within one doubling dilution was 99.4%.

**Figure 2:**
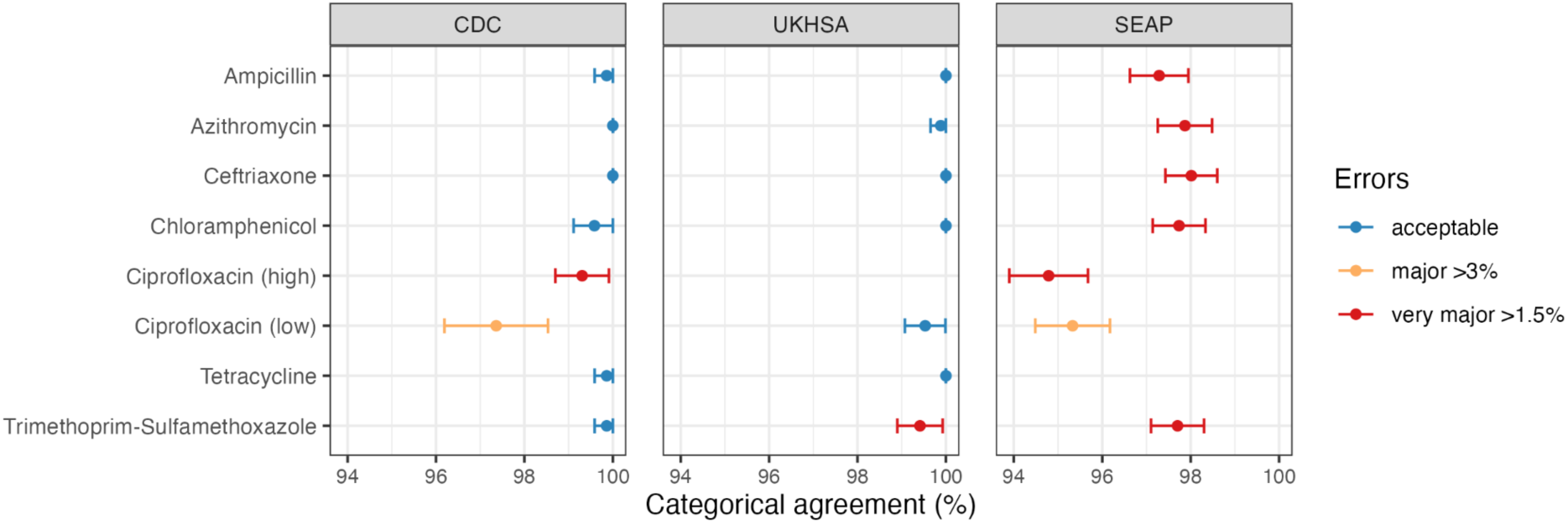
Validation of Typhi Mykrobe AMR predictions vs AMR phenotypes Summary of Typhi Mykrobe’s genotype-based clinical resistance categorisations compared with susceptibility phenotypes, for 4,018 isolates with publicly available matched genome and phenotype data. Comparisons are summarised as categorical agreement, and calculated separately for the three different source datasets, which used different phenotyping methods and interpretive standards (US CDC: MIC, CLSI; UKHSA: MIC, EUCAST; SEAP: disk diffusion, EUCAST; see **Methods**). Major errors were defined as susceptible isolates with AMR determinants detected (reported as R by Mykrobe). Very major errors were defined as resistant isolates with no AMR determinants detected (reported as S by Mykrobe). Error bars are shown that represent 95% confidence interval of proportion. Note UKHSA tested amoxicillin rather than ampicillin, the result is reported in the ampicillin row.

**Figure 3:**
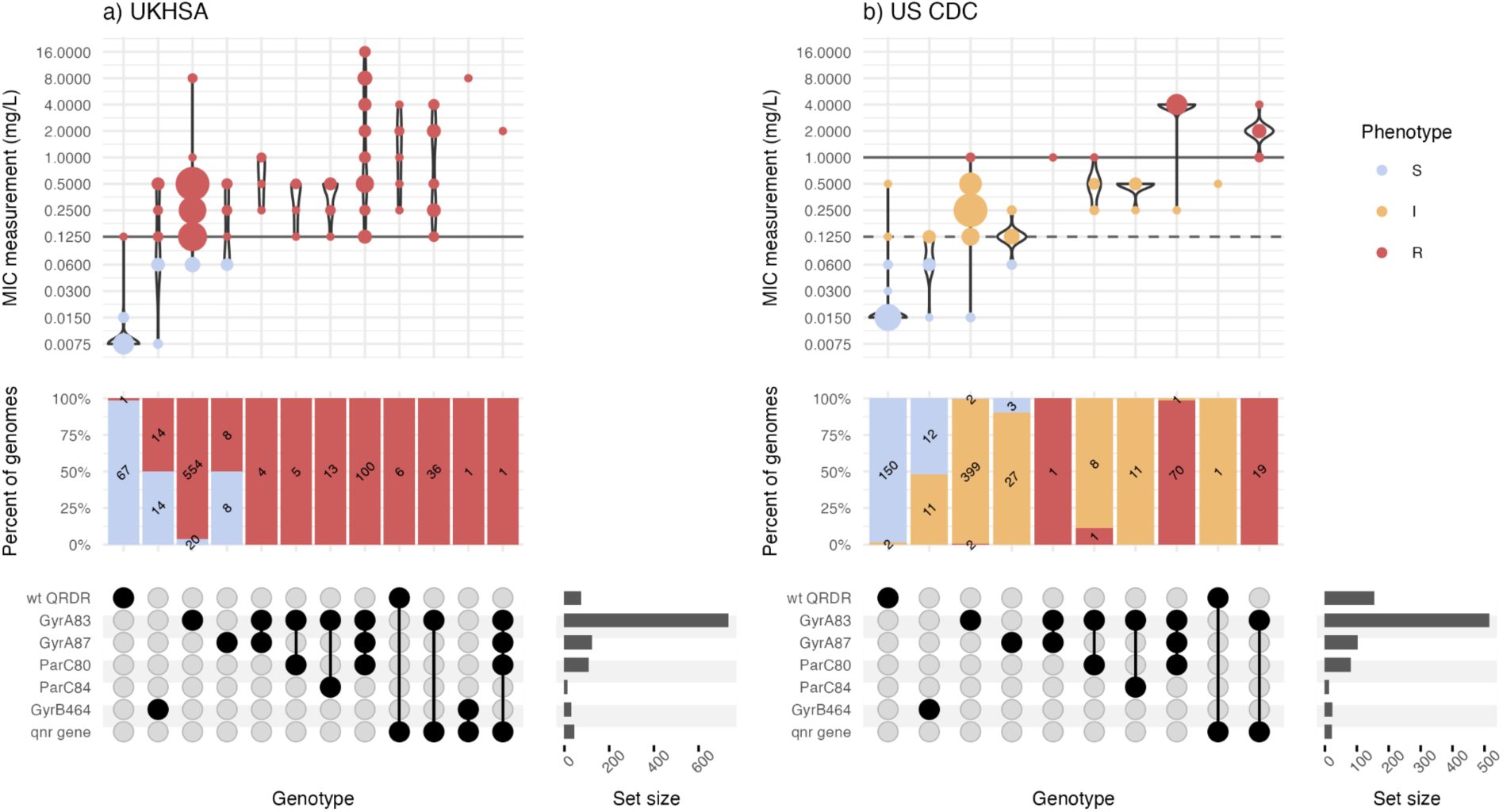
Typhi Mykrobe genotypes vs minimum inhibitory concentration from two reference laboratories, for ciprofloxacin. **a)** UKHSA dataset, using EUCAST. **b)** US CDC dataset, using CLSI. Each column represents the set of isolates in which Typhi Mykrobe identified a unique combination of genetic determinants, indicated in the panel at the bottom. For each column, the violin plots show the distribution of ciprofloxacin MIC values, and stacked barplots show the proportion of genomes called as S, I, or R (coloured as per inset legend, and labelled with counts). The solid horizontal lines on the violin plots mark the R breakpoint, the dashed line marks the CLSI I breakpoint used for the US CDC dataset.

Major errors were infrequent (<3% prevalence) for all drugs in all datasets (**Figure 2**), except for ciprofloxacin (**Supplementary Table 5**). Most of the ciprofloxacin major errors were in isolates whose MIC or disk zone measurement were close to the breakpoint (**Figures 3, 4c**); all carried well-characterized molecular resistance determinants (QRDR mutations and/or *qnr* genes, see **Figures 3, 4c**), and in all but one case the same determinants were also detected in the corresponding genome assemblies by Pathogenwatch (details below).

Very major errors (resistant isolates with no AMR determinants detected, thus reported as S by Mykrobe) were also infrequent (<1.5% prevalence) for the reference-laboratory MIC datasets, but errors were higher in the disk diffusion dataset (**Figure 2**). The only drugs showing major or very major errors compared with the reference-laboratory MIC data were ciprofloxacin and trimethoprim-sulfamethoxazole (**Figure 2**). For ciprofloxacin, the US CDC categorisation of high-level resistance (using the CLSI ‘R’ threshold, ≥1 mg/L) yielded four very major errors (R isolate called as ‘I’ by Mykrobe). Two had a single mutation detected (GyrA-S83F) and two had double QRDR mutations (GyrA-S83F plus GyrA-D87N or ParC-S80I) (see **Figure 3**); Pathogenwatch calls agreed in all instances. Importantly, all four isolates had MIC=1 mg/L, i.e. they agreed with the genotype-based call within one doubling dilution. UKHSA uses the EUCAST standards, which have a single (low-level) breakpoint for ciprofloxacin (R >0.06 mg/L), and this yielded just two major errors (2.9%) and two very major errors (0.26%) (see **Figure 3**).

For trimethoprim-sulfamethoxazole, Mykrobe analysis of the US CDC dataset yielded a single error (R isolate called as S) but the MIC was 4 mg/L, i.e. within one doubling dilution, and the isolate had no *sul* or *dfr* genes that were identified by Mykrobe or Pathogenwatch. Mykrobe analysis of the UKHSA dataset yielded five errors for trimethoprim-sulfamethoxazole; all were R isolates called as S by Mykrobe, which detected *sul* genes but no *dfrA* genes. Interestingly, all five isolates tested susceptible to trimethoprim alone (MIC ≤0.5, well below the >4 mg/L breakpoint), supporting the absence of *dfrA* genes; these were also not detected by Pathogenwatch analysis of the corresponding genome assemblies, or by UKHSA’s internal pipeline [44]. Further investigation of these genome assemblies confirmed the presence of wild-type *folA* and *folP*, and no additional determinants were reported by CARD RGI (v6.0.3).

Mykrobe analysis of the SEAP dataset showed lower overall categorical agreement with AST phenotypes (95.3–98.2%). As the testing was done using disk diffusion, it was not possible to assess the categorical agreement within one doubling dilution of a MIC. However, most of the errors were within 2–4 mm of the breakpoint (see **Figure 4**), and in all but one instance, the Mykrobe genotype result matched that called by Pathogenwatch from the corresponding genome assemblies. The only exception was a ciprofloxacin resistant isolate (ERR4325960), which Mykrobe reported as carrying GyrA*-*S83F with no other determinants, but Pathogenwatch detected both GyrA*-*S83F and *qnrS* in the assembly.

**Figure 4:**
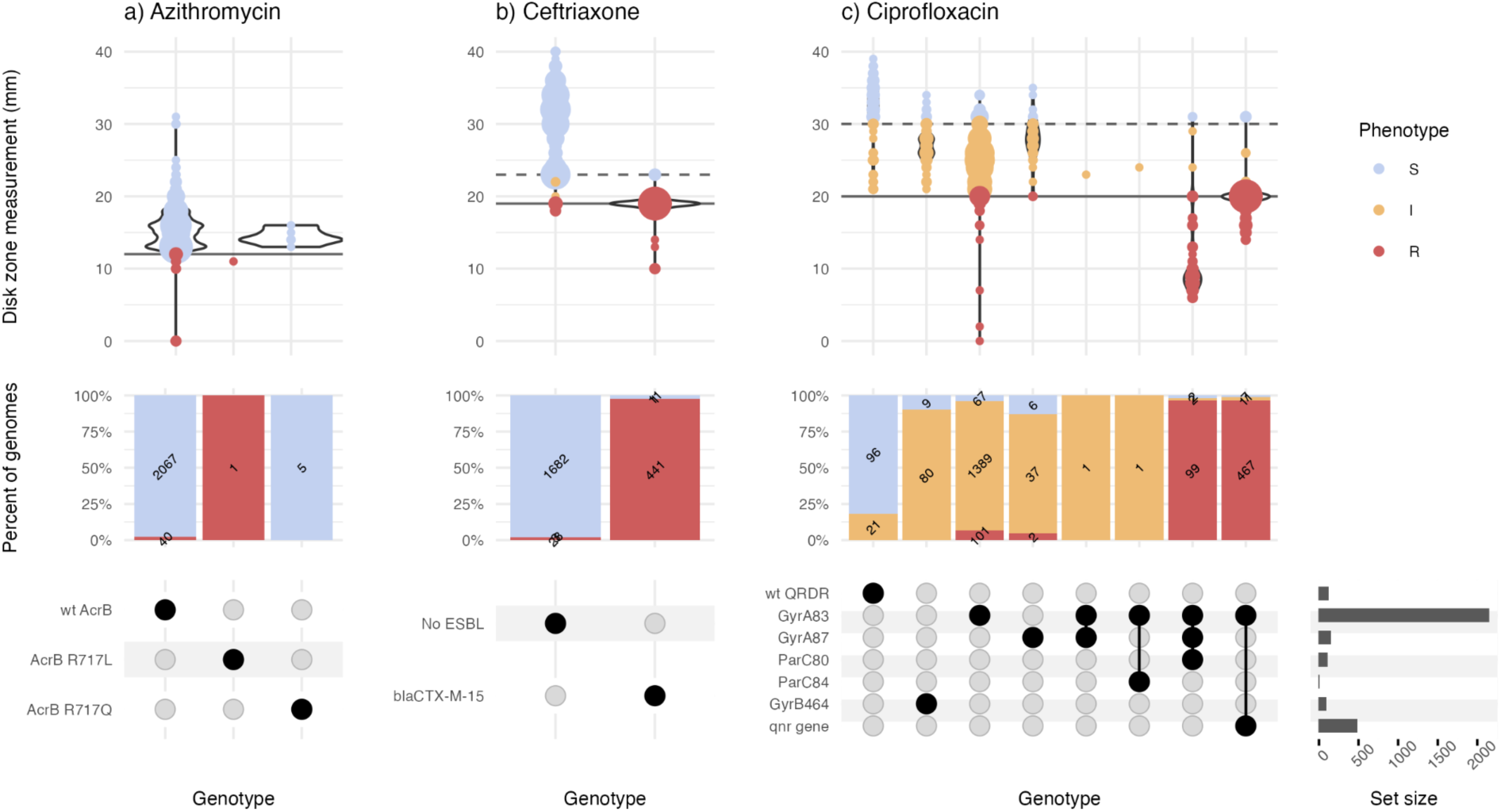
Typhi Mykrobe genotypes vs disk diffusion data from the SEAP study, for three drug classes Data for three drugs are shown in panels **a)** Azithromycin, **b)** Ceftriaxone, and **c)** Ciprofloxacin. Each column represents the set of isolates in which Typhi Mykrobe identified a unique combination of genetic determinants, indicated in the panel at the bottom. For each column, the violin plots show the distribution of disk diffusion values for azithromycin, ceftriaxone or ciprofloxacin. The stacked barplots show the proportion of genomes called as S, I, or R (coloured as per inset legend, and labelled with counts). The solid horizontal lines on the violin plots mark the R EUCAST breakpoint used for the SEAP disk diffusion data. The dashed line marks the EUCAST breakpoint for I.

### Validation on nanopore reads

We explored the accuracy of Typhi Mykrobe for genotyping ONT long-read data by comparing results from 98 pairs of ONT to Illumina reads generated from the same isolates. Data were contributed by members of the Global Typhoid Genomics Consortium and generated using different library preparation methods and sequencing devices (see **Supplementary Table 6**). Mykrobe called one Illumina read set and five ONT read sets as non-Typhi (mean ONT read depths estimated as 0.04×, 4.06×, 8.01×, 26.40×, 27.85×, all sequenced on R9 flowcells), these were excluded from analysis of genotype accuracy as no genotype is called on isolates identified as non-Typhi.

Among the n=92 Typhi genomes called from both ONT and Illumina reads, the Typhi Mykrobe genotype lineage calls matched for n=89 (96.7%). For all three discrepant isolates, (all sequenced on R.9 or R.9.4 flowcells), the set of SNV markers detected from ONT reads closely matched those detected from Illumina reads, but with one or two markers missing from the ONT-based calls and no unexpected markers reported resulting in final genotype calls of 4.3.1.1 vs 4.3.1, 4.3.1.2 vs 4.3.1, 2.1.7.2 vs 2.1.7. For the missing markers, Mykrobe reported low and variable ONT read depths, with ranges 3–9, 4–31 and 1–14, suggesting that the errors are likely due to lack of informative ONT reads at the marker loci. Notably, the overall ONT read depths for these were reasonable (mean 35×, 170× and 90× respectively), suggesting the lack of informative reads for genotyping may be caused by contamination from non-Typhi reads, or high error rates resulting in low rates of *k*-mer matching.

In this study, the five ONT genomes that had ‘not Typhi’ calls were further explored. These were all classified as *Salmonella enterica*. However, Typhi was not the top serovar from read-level classification, instead *Salmonella* Infantis was the top serovar for BZD62J, while *Salmonella* Typhimurium was the top serovar for the remaining four. Further, the presence of the marker genes of *invA* and the seven MLST genes were detected in the genomes at read depths consistent with the genome mean read depth reported in **Supplementary Table 6**, with the gene *invA* was detected in three of the five genomes from Blast analysis of the Flye assemblies. The gene *invA* was not detected in the 611427 ONT assembly and only had a read depth of 0.43×, and BZD62J failed to assemble due to insufficient reads and was found to have no reads mapping to *invA.* MLST analysis of the three complete genomes with *invA* detected reported, were found to have partial matches to known alleles for all seven MLST genes and no reported MLST profile. Of note, these five genomes had low read depth (0.04–28×), sequencing on older R9 flowcells, and basecalling with older versions of guppy (**Supplementary Table 6**). The failure of Typhi Mykrobe to identify these as Typhi is likely due to a combination of potential contamination, low read depth and high error rates resulted in low rates of *k*-mer matching to the probes. This highlights the importance of QC of the read data, as Typhi Mykrobe provides some metrics for confidence of the calls, but does not itself perform detailed QC.

AMR marker detection from ONT reads was highly concordant with Illumina-based calls, with 98.77% overall agreement across the n=736 isolate-drug combinations (92 isolates across 8 drugs; see **Supplementary Table 7**). There were nine genomes with disagreement of AMR calls; five of these had a marker detected in Illumina reads but not ONT reads (n=1 *catA1*, n=1 *qnr,* n=3 QRDR mutations), three had GyrB*-*464F detected in ONT reads but not Illumina, and one had GyrB*-*S464F detected in Illumina reads and GyrA*-*S83F in ONT. In all nine cases, the Typhi Mykrobe call from Illumina reads matched that from Pathogenwatch analysis of the corresponding assemblies of Illumina reads. There was no overlap between ONT read sets with AMR discrepancies and those with lineage genotype discrepancies. Overall, ONT-based resistance prediction showed complete agreement with Illumina for six of the eight drugs, 98.9% agreement for chloramphenicol resistance, and 91% for ciprofloxacin resistance.

Plasmid marker detection from ONT data was also highly concordant with Illumina-based calls, with 99.67% overall agreement across the n=1,196 isolate-marker combinations (92 isolates across 13 markers; see **Supplementary Table 8**). Four genomes had discrepant plasmid marker calls, all with a marker detected from Illumina reads but not from ONT reads. One isolate (DRR070993) had a typical IncHI1 MDR plasmid profile reported from Illumina reads (IncHI1A, IncHI1BR27, IncHI1_ST6 indicating presence of IncHI1-ST6 plasmid, with *catA1*, *dfrA7*, *bla*_TEM-1_, *sul1*, *sul2*), but the ONT read profile was missing the *IncHI1A* and *catA1* markers (and missing *catA1* in the corresponding Illumina assembly and a mismatch for *IncHI1A*). The other three isolates had pHCM2 detected in the Illumina reads and in the corresponding assemblies [14] but not in ONT reads; for these isolates the ONT and Illumina sequencing was done on different DNA extracts so this may reflect plasmid loss in the culture used for ONT sequencing.

## Discussion

Typhi Mykrobe provides detection and genotyping of key features of clinical and public health importance, including assigning lineages, AMR determinants, and plasmid replicons (**Figure 1**). Compared with other tools for Typhi genotyping, Mykrobe has the advantage of not requiring any pre-processing, aligning, or assembling of reads, but instead takes as inputs raw FASTQ files. This simplifies bioinformatics workflows and reduces time-to-result as Typhi Mykrobe can return a full genotype profile in typically <1 minute. Importantly, although Typhi Mykrobe is not a QC tool *per se*, it provides useful data on the read-level support for individual genotype markers, which can be used to help identify potentially mixed samples. As a command-line tool that is easy to install locally using Bioconda, Typhi Mykrobe is suitable for research and public health laboratory settings, facilitates standardized reporting of Typhi genotypes without the need to share sequence data, and enables broader access to bioinformatics functionality across a variety of settings.

Typhi Mykrobe showed very high genotyping accuracy, for GenoTyphi lineage assignment (**Table 1**), and detection of AMR and plasmid markers (**Tables 2-3**), compared with existing genotyping tools. Of note, the included IncY, pO111 and IncFIB for the cryptic pHCM2 replicon targets have recently been shown to be sometimes carried on phage-plasmids [56,57], in addition to conjugative plasmids such as the IncY plasmids associated with XDR Typhi [26].

Future work is required to identify additional targets to delineate the phage-plasmids, and the Typhi Mykrobe software can be readily updated to include new markers.

In addition, we show that the panel of AMR markers currently included in Typhi Mykrobe can be used to predict AMR phenotypes with high accuracy, demonstrating very high concordance with S/I/R calls made from reference-laboratory MICs (**Figure 2**). Concordance was slightly lower when compared with research laboratory disk diffusion data. However, as the phenotypic resistance prevalence rates were similar across the three datasets, it is reasonable to assume that the higher error rates in the disk diffusion dataset compared with the two reference-laboratory MIC datasets most likely reflect comparatively lower accuracy of the disk diffusion phenotype measurements, which are known to yield lower accuracy for some drugs such as azithromycin [58,59] and in this case were completed across multiple laboratories. There are currently no formal standards for assessing the accuracy of AMR phenotype predictions from genomic data[60]; however, applying the standards for FDA licensing of new AST devices, Typhi Mykrobe easily passed the target threshold for categorical agreement (>99% agreement within one doubling dilution for reference-laboratory MIC data, >95% agreement for disk-diffusion data; vs >90% target) (**Figure 2**).

Typhi Mykrobe also showed very low rates of major errors and very major errors compared with the reference-laboratory MIC data, and for nearly all drugs these fell below the acceptable thresholds for a new AST device (<3% and <1.5%, respectively) (**Figure 2**). The exception was trimethoprim-sulfamethoxazole, for which five UKHSA isolates tested resistant but Typhi Mykrobe predicted as susceptible due to a lack of *dfrA* genes, which are thought to be required for resistance to trimethoprim-sulfamethoxazole along with *sul* genes, which were detected in these genomes. Other genotyping tools (Typhi Pathogenwatch, CARD RGI) also could not identify any known mechanisms of resistance to trimethoprim or trimethoprim-sulfamethoxazole in these genomes, and the isolates tested sensitive to trimethoprim. Therefore, we hypothesise these isolates may harbour efflux mutations or a novel mechanism of resistance to trimethoprim-sulfamethoxazole that does not result in resistance to trimethoprim alone. Further analyses to determine the exact resistance mechanism was beyond the scope of this study.

Ciprofloxacin showed very low error rates in the UKHSA MIC data, interpreted using the single low-level resistance threshold (MIC >0.06 mg/L) recommended by EUCAST, but slightly higher error rates for the US CDC MIC data, which was interpreted using low-level and high-level resistance thresholds (although still below the acceptable thresholds within one doubling dilution). Carbapenemase genes were added to the Typhi Mykrobe panel, but this resistance is newly emerging and there was only a single carbapenem-resistant isolate sequenced at the time of testing (in which *bla*NDM-5 was correctly identified by Typhi Mykrobe), therefore we could not yet assess accuracy of phenotype prediction for carbapenems.

Finally, we demonstrated that Typhi Mykrobe can be used to genotype ONT reads, showing high concordance with results obtained from the analysis of Illumina reads generated from the same isolates (96.74% agreement on GenoTyphi lineage calls, 98.77% on AMR markers and 99.67% on plasmid markers). We did not have suitable data to directly assess ONT-based AMR predictions with AST data. ONT-based and Illumina-based drug-level predictions of AST phenotypes showed high agreement; however, future studies assessing ONT data compared to reference-laboratory MIC data would be useful to confirm the accuracy of ONT data for phenotype prediction. As one might expect, many of the errors occurred in isolates with lower read depth. However, as the matched ONT-Illumina data available for testing were contributed by different laboratories that used different ONT protocols and devices for library preparation, sequencing, and base calling, we were unable to perform a detailed assessment of the impact of read depth and quality on genotyping performance. The accuracy of ONT reads has greatly increased in recent years and is approaching Illumina read accuracy of ∼99%; hence modern ONT data would be expected to perform similarly to Illumina data [61] (doi.org/10.5281/zenodo.10397817). Future studies assessing the impact of read depth and base calling methods on the performance of Typhi Mykrobe on ONT data would be desirable to help guide laboratories on the most appropriate methods and target yield for ONT experiments aiming to produce reliable genomic surveillance data for Typhi.

## Conclusions

Typhi Mykrobe provides rapid and accurate genotyping of Typhi genomes directly from Illumina reads, including (i) assignment of GenoTyphi lineage; (ii) detection of AMR determinants and prediction of corresponding AMR phenotypes demonstrating >99% categorical agreement with reference-laboratory MIC data; and (iii) detection of plasmid replicons. The software also performs well on direct genotyping from ONT reads information, although further investigation is needed to assess the accuracy of AMR phenotype prediction.

## List of abbreviations

AMR: antimicrobial resistance
AST: antimicrobial susceptibility testing
CARD: Comprehensive Antibiotic Resistance Database
CLSI: Clinical and Laboratory Standards Institute
EUCAST: The European Committee on Antimicrobial Susceptibility
Testing FDA: United States Food and Drug Administration
MDR: multidrug resistance
MIC: minimum inhibitory concentration
ONT: Oxford Nanopore Technologies
QRDR: quinolone resistance-determining regions
SEAP: Surveillance for Enteric Fever in Asia Project
SNV: single nucleotide variant
UKHSA: United Kingdom Health Security Agency
US CDC: United States of America Centers for Disease Control and Prevention
WGS: whole genome sequencing
XDR: extensively drug resistant

## Other sections

## Availability and requirements

Project name: GenoTyphi

Project home page: https://github.com/typhoidgenomics/genotyphi Operating system(s): Platform independent

Programming language: Python3 or higher Other requirements: Mykrobe

License: GNU General Public License

Any restrictions to use by non-academics: None

Project name: Mykrobe

Project home page: https://github.com/Mykrobe-tools/mykrobe Operating system(s): Platform independent

Programming language: Python3 or higher

Other requirements: anytree 2.8.0 or higher, Biopython 1.79 or higher, PyVCF3 1.0.3 or higher, requests 2.27.1 or higher, mongoengine 0.24.1 or higher, cython 0.29.28 or higher, numpy 1.22.0 or higher, pytest License: MIT License

Any restrictions to use by non-academics: None.

## Declarations

## Ethics approval and consent to participate

Each contributing study or surveillance program obtained local ethical and governance approvals, as reported in the primary publication for each dataset. Inclusion of data that were not yet public domain by August 2021 was approved by the Observational / Interventions

Research Ethics Committee of the London School of Hygiene and Tropical Medicine (ref #26408), on the basis of details provided on the local ethical approvals for sample and data collection (listed in Carey et al, 2023, Supplementary file 1).

## Consent for publication

Not applicable

## Availability of data and materials

All code and instructions for genotyping isolates with GenoTyphi using Mykrobe is available in the GitHub repository at https://github.com/typhoidgenomics/genotyphi. The version reported in this paper is v2.1 (DOI: <u>10.5281/zenodo.13859721</u>)

The datasets and analysis code used in this article are available in a GitHub repository at https://github.com/typhoidgenomics/TyphoidGenomicsConsortiumMykrobe.

## Authors’ Contributions

DJI, JH, ZAD and KEH – Conceptualization. DJI, JAK, ZAD and KEH - Data curation.

JH, MH, ZI, ZAD and KEH – Software

DJI, JH, RRW, ZAD and KEH - Methodology

DJI, JK, AOA, NA, SA, PMA, IIB, MEC, MAC, JAC, PDG, HI, JJJ, AK, KHK, MM, SN, SO, INO, SIAR, AR, DR, VS, MJS, AMS, GTS, KAT, RRW, ZAD and KEH - Formal analysis

DJI, JH, AOA, NA, SA, PMA, IIB, MC, MAC, JAC, PDG, BPH, HI, JJJ, LMJ, AK, KHK, JK, MML, MM, SN, SO, INO, PEO, SIAR, AR, DAR, VS, MJS, AMS, GTS, KAT, RRW, ZAD and KEH - Investigation, DJI and KEH - Visualization, DJI, ZAD and KEH - Writing – original draft, DJI, JH, ZI, MH, JAK, AOA, NA, SA, PMA, IIB, MEC, MAC, JC, PDG, BPH, HI, JJJ, LMJ, AK, KHK, JK, ML, MM, SN, SO, INO, PEO, SIAR, AR, DR, VS, MJS, AMS, GTS, KT, RRW, ZAD and KEH - Writing – review and editing

All authors read and approved the final manuscript.

## Acknowledgements

MAC and SN are affiliated to the National Institute for Health Research Health Protection Research Unit (NIHR HPRU) in Genomics and Enabling Data at University of Warwick in partnership with the UK Health Security Agency (UKHSA). MAC and SN are based at UKHSA. MH is affiliated to the NIHR Health Protection Research Unit in Healthcare Associated Infections and Antimicrobial Resistance at Oxford University. The views expressed are those of the author(s) and not necessarily those of the NIHR, the Department of Health and Social Care or the UK Health Security Agency.

SIAR from the icddr,b is grateful to the Governments of Bangladesh, and Canada for providing unrestricted support.

NA is grateful for support from ICMR India (IIRP-2023-2500).

KAT thanks the epidemiology and laboratory partners in state and local health departments. The findings and conclusions in this report are those of the authors and do not necessarily represent the official position of the Centers for Disease Control and Prevention. Names of specific vendors, manufacturers, or products are included for public health and informational purposes; inclusion does not imply endorsement of the vendors, manufacturers, or products by the Centers for Disease Control and Prevention or the US Department of Health and Human Services.

AMS thanks all participants of the GERMS-SA Laboratory Surveillance Network, South Africa, for submission of clinical isolates of Salmonella species to the National Institute for Communicable Diseases; the surveillance network includes laboratories belonging to the Department of Health (NHLS laboratories) and laboratories that form part of the private sector. We thank all the lab members involved in SEFI reference lab activities at CMC Vellore implicated in culture maintenance and ONT data generation.

This work was supported by Monash eResearch capabilities, including the high performance computer M3 and Research Data Storage.

## Global Typhoid Genomics Consortium Group Authorship

Ali H. Abbas^1^, Safina Abdul Razzak^2^, Mabel K. Aworh^3^, Stephen Baker^4^, Buddha Basnyat^5^, Ka Lip Chew^6^, Thomas C. Darton^7^, Andrew R. Greenhill^8^, Madhu Gupta^9^, Mochammad Hatta^10^, Aamer Ikram^11^, Dasaratha Ramaiah Jinka^12^, A. C. Lauer^13^, Octavie Lunguya^14^, Jaspreet Mahindroo^15^, Tapfumanei Mashe^16^, Elli Mylona^17^, Maria Pardos de la Gandara^18^, Andrew J. Pollard^19^, Saikt Rahman^20^, Elrashdy M Redwan^21,22,23^, Jean Pierre Rutanga^24^, Jivan Shakya^25^, Arif Tanmoy^26^, Mathew S. Thomas^27^, Sandra Van Puyvelde^28^, Jacqueline M Wright^29^

1 Department of Microbiology, Faculty of Veterinary Medicine, University of Kufa, Kufa-Najaf Street, Najaf, Iraq

2 Department of Paediatrics and Child Health, Aga Khan University, P.O. Box 3500, National Stadium Rd, Karachi, Sindh, Pakistan.

3 North Carolina State University, 1060 William Moore Drive, Raleigh, NC 27607, USA

4 University of Cambridge, JCBC, Puddicome way, Cambridge, CB2 0AW, United Kingdom

5 OUCRU-Nepal, Jhamshikhel, Lalitpur, Nepal

6 National University Hospital, 5 Lower Kent Ridge Road, Singapore 119074, Republic of Singapore

7 Clinical Infection Research Group, School of Medicine and Population Health, University of Sheffield, Western Bank, Sheffield, S10 2TN, United Kingdom

8 Federation University Australia, Gippsland Campus, Churchill, Victoria, Australia

9 Postgraduate Institute of Medical Education and Research, sector 12, Chandigarh. CMC Vellore, India

10 Department of Molecular Biology and Immunology, Faculty of Medicine, Hasanuddin University, Makassar, 90245, Indonesia

11 Fauji Foundation, Rawal Road, Rawalpindi, Pakistan

12 RDT Hospital, Bathalapalli, Srisathya Sai District, 515661, India

13 Centers for Disease Control and Prevention, Atlanta, United States of America

14 Department of Microbiology, Institut National de Recherche Biomédicale, Kinshasa, Democratic Republic of the Congo

15 MRC Centre for Global Infectious Disease Analysis, School of Public Health, Imperial College London, London, W12 0BZ, United Kingdom

16 World Health Organisation, Zimbabwe

17 Cambridge Institute of Therapeutic Immunology and Infectious Disease (CITIID), University of Cambridge, Cambridge CB1 0AW, United Kingdom

18 Institut Pasteur, France

19 Oxford Vaccine Group, Department of Paediatrics, University of Oxford and the NIHR Oxford Biomedical Research Centre, Oxford, UK

20 RDT Hospital, Bathalapalli, Srisathya Sai District, 515661, India

21 Department of Biological Science, Faculty of Science, King Abdulaziz University, Jeddah, Saudi Arabia

22 Centre of Excellence in Bionanoscience Research, King Abdulaziz University, Jeddah 21589, Saudi Arabia

23 Therapeutic and Protective Proteins Laboratory, Protein Research Department, Genetic Engineering and Biotechnology Research Institute, City of Scientific Research and Technological Applications, New Borg EL-Arab 21934, Alexandria, Egypt.

24 University of Rwanda, KK 737 street, Kigali, B.P 3900, Rwanda

25 Institute for Research in Science and Technology, Sachal-2, Lalitpur, Nepal

26 Child Health Research Foundation, Dhaka, Bangladesh

27 CMAI, ICMDA

28 University of Antwerp, Antwerp, Belgium

29 Institute of Environmental Sciences and Research 27 Creyke Road, Christchurch, 8041, New Zealand

