## Supplementary_materials for "Typhi Mykrobe: fast and accurate lineage identification and antimicrobial resistance genotyping directly from sequence reads for the typhoid fever agent *Salmonella* Typhi"

Ingle et al.

Supplementary tables, and code to generate tables and figures, is in the Typhoid Genomics Consortium Typhi Mykrobe github:
https://github.com/typhoidgenomics/TyphoidGenomicsConsortiumMykrobe

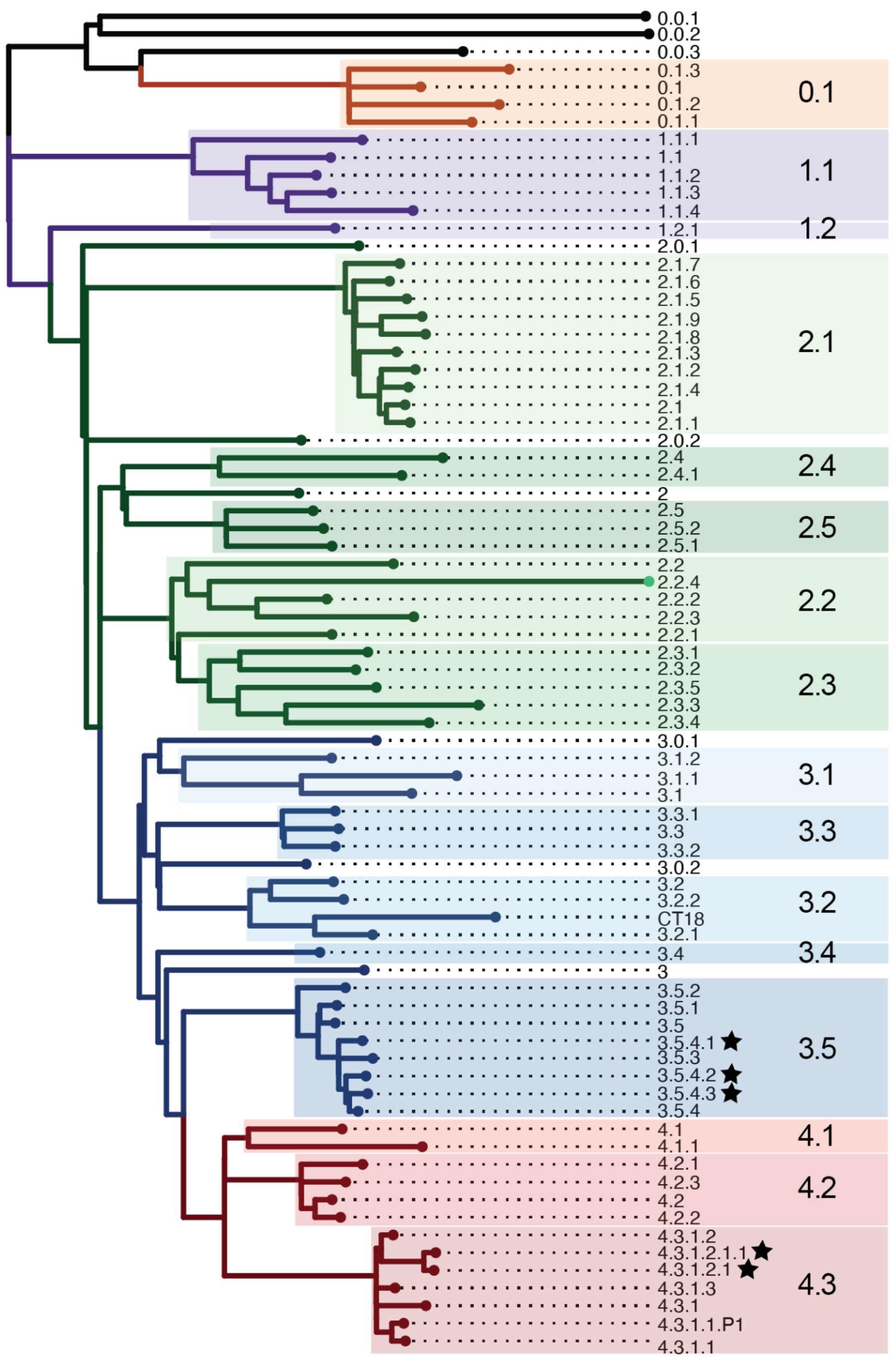


**Supplementary Figure 1: Overview of the GenoTyphi scheme**
Phylogenetic tree backbone showing the relationships between the lineages, clades and subclades. Tree tips represent unique genotypes as labeled, and background shading highlights clades (labeled in larger font). The black stars indicate genotypes added to the scheme in the 2022 Technical Report (https://zenodo.org/doi/10.5281/zenodo.7407984 )

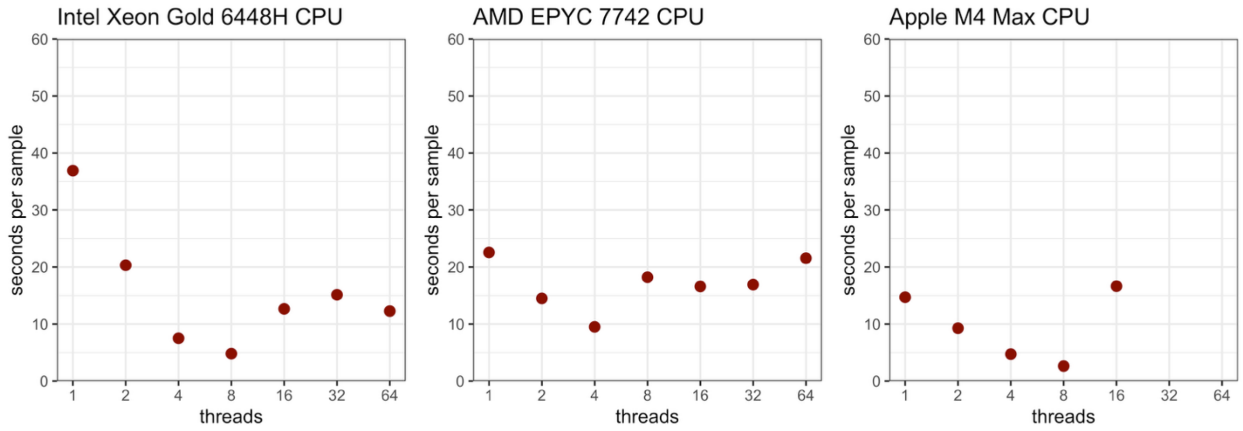


**Supplementary Figure 2: Average run-time of Typhi Mykrobe of 100 Typhi genomes**
The average run time for 100 Typhi genomes on three different computers. The number of threads used is shown on the x-axis. The average time (seconds per sample) to run on each genome is shown on the y axis.

**Supplementary Table 1.** Tabulated Typhi Mykrobe output table for all genomes included in validation analyses (available in github).

**Supplementary Table 2:** Genome data used for validation (available in github).

**Supplementary Table 3.** Details of AMR genotype calls comparison (available in github).

**Supplementary Table 4.** Genome data for all isolates with publicly available antimicrobial susceptibility testing (AST) (available in github).

**Supplementary Table 5.** Comparison and error rates for AMR genotype and phenotype data (available in github).

**Supplementary Table 6.** Details of validation of typing from nanopore reads (available in github).

### Supplementary Table 7. Validation of AMR genotyping from ONT reads (vs Illumina)

| Drug | Illumina | ONT (vs Illumina) | N | Agreement |
|---|---|---|---|---|
| Ampicillin | *bla*TEM-1 | agree | 35 | 100% |
| | no marker | agree | 57 | |
| Azithromycin | no marker | agree | 92 | 100% |
| Ceftriaxone | *bla*CTX-M-15 | agree | 5 | 100% |
| | *bla*SHV-12 | agree | 1 | |
| | no marker | agree | 86 | |
| Chloramphenicol | *catA1* | agree | 25 | 98.99% |
| | | *no marker | 1 | |
| | no marker | agree | 66 | |
| Ciprofloxacin | 1 QRDR | agree | 34 | 91.30% |
| | | *different QRDR | 1 | |
| | | *no marker | 3 | |
| | 2 QRDR | agree | 2 | |
| | 3 QRDR | agree | 3 | |
| | 1 QRDR + *qnr* | agree | 5 | |
| | | *1 QRDR | 1 | |
| | no marker | agree | 40 | |
| | | *1 QRDR | 3 | |
| Sulfonamides | *sul1* | agree | 5 | 100% |
| | *sul2* | agree | 9 | |
| | *sul1;sul2* | agree | 24 | |
| | no marker | agree | 54 | |
| Trimethoprim | *dfrA7* | agree | 26 | 100% |
| | *dfrA14* | agree | 4 | |
| | *dfrA15* | agree | 2 | |
| | no marker | agree | 60 | |
| Tetracycline | *tetA(A)* | agree | 7 | 100% |
| | *tetA(B)* | agree | 2 | |
| | no marker | agree | 83 | |

**Supplementary Table 8. Validation of plasmid marker detection from ONT reads (vs Illumina)**

| Rep marker | Illumina | ONT (vs Illumina) | N | Agreement |
|---|---|---|---|---|
| IncFIAHI1 | present | agree | 1 | 100% |
| | absent | agree | 91 | |
| IncHI1A | present | agree | 1 | 98.91% |
| | present | *absent | 1 | |
| | absent | agree | 90 | |
| IncHI1BR27 | present | agree | 2 | 100% |
| | absent | agree | 90 | |
| IncHI1_ST6 | present | agree | 2 | 100% |
| | absent | agree | 90 | |
| IncHI2A | absent | agree | 92 | 100% |
| IncY | present | agree | 7 | 100% |
| | absent | agree | 85 | |
| IncX3 | present | agree | 1 | 100% |
| | absent | agree | 91 | |
| IncI1 | absent | agree | 92 | 100% |
| IncL_M | absent | agree | 92 | 100% |
| IncFIB_pHCM2 | present | agree | 12 | 96.74% |
| | present | *absent | 3 | |
| | absent | agree | 77 | |
| IncFIB_K | present | agree | 2 | 100% |
| | absent | agree | 90 | |
| IncN | present | agree | 3 | 100% |
| | absent | agree | 89 | |
| z66 | absent | agree | 92 | 100% |
| **Total** | present | agree | 31 | **99.67%** |
| | present | *absent | 4 | |
| | absent | agree | 1161 | |
